# Distinct alterations in white matter properties and organization related to maternal treatment initiation in neonates exposed to HIV but uninfected

**DOI:** 10.1101/2024.01.11.575169

**Authors:** Ndivhuwo Magondo, Ernesta M. Meintjes, Fleur L. Warton, Francesca Little, Andre J.W. van der Kouwe, Barbara Laughton, Marcin Jankiewicz, Martha J. Holmes

## Abstract

HIV exposed-uninfected (HEU) infants and children are at risk of developmental delays as compared to uninfected unexposed (HUU) populations. The effects of exposure to *in utero* HIV and ART regimens on the HEU the developing brain are not well understood.

In a cohort of 2-week-old newborns, we used diffusion tensor imaging (DTI) tractography and graph theory to examine the influence of HIV and ART exposure *in utero* on neonate white matter integrity and organisation. The cohort included HEU infants born to mothers who started ART before conception (HEU_pre_) and after conception (HEU_post_), as well as HUU infants from the same community. We investigated HIV exposure and ART duration group differences in DTI metrics (fractional anisotropy (FA) and mean diffusivity (MD)) and graph measures across white matter.

We found increased MD in white matter connections involving the thalamus and limbic system in the HEU_pre_ group compared to HUU. We further identified reduced nodal efficiency in the basal ganglia. Within the HEU_post_ group, we observed reduced FA in cortical-subcortical and cerebellar connections as well as decreased transitivity in the hindbrain area compared to HUU.

Overall, our analysis demonstrated distinct alterations in white matter integrity related to the timing of maternal ART initiation that influence regional brain network properties.

## Introduction

The developing brain is at its most vulnerable *in utero*, where it depends on the maternal environment to provide nutrients and protection. Disruptions to the maternal ecosystem may occur as a result of viral, bacterial or parasitic infections. Studies have established that even indirect contact with a viral infection *in utero* can lead to adverse effects in fetuses, suggesting that the maternal immune response and symptoms of infections may contribute to adverse prenatal development^1,2^. While treatments for infection that may restore maternal health are available, such as antiviral and antibiotics, there may be negative neurodevelopmental consequences for the fetus^3,4^.

HIV (human immunodeficiency virus) targets and alters the immune system. In women living with HIV, the virus can adversely affect maternal health as well as fetal development. During pregnancy, antiretroviral therapy (ART) provides protection to the fetus by preventing HIV proliferation in the mother, allowing her immune system to strengthen. Due to increased availability of ART, the rate of vertical HIV transmission has decreased substantially leading to a growing population of HIV exposed uninfected (HEU) infants. Although uninfected, HEU infants demonstrate higher risks of cognitive deficits, particularly in language and motor skills, compared to HIV unexposed uninfected (HUU) populations^5^.

White matter connections play an integral role in cognition because they facilitate rapid transmission of information between different brain areas^6^. In utero, white matter organization primarily involves the development of axonal fibers. Axonal pathways start developing at 8 post conceptual weeks, the end of the embryonic stage, into the neonatal period. Pre-myelination begins in the second trimester as fibers mature and myelin forming cells develop. Mature myelin appears between 20- and 28 weeks of gestational age (GA), with the portion of total brain volume that contains myelinated white matter increasing from 1 to 5% between 36- and 40 weeks GA^7^. As a result, disruptions to the maternal environment across the stages of pregnancy can disrupt different aspects of developing white matter.

The basic structural and functional wiring of the brain is already in place at birth^6^. During the last trimester of pregnancy, short and long connections between brain regions develop to create an early adult-like organization of neural networks^8^. At birth, a large network of connections is already present, which may be referred to as a connectome^9^. Given the time sensitive nature of white matter development, the timing of ART initiation during pregnancy may contribute to its ability to provide protection or disruption of white matter development.

Diffusion tensor imaging (DTI) yields quantitative measures reflecting properties of white matter. Biological models of developing white matter *in utero* and early infancy based on imaging measures have been proposed from which typical white matter development, axon growth/development, pre-myelination and myelination, can be inferred based on changes in DTI parameters^10^.

While several publications have looked at the effects of HIV exposure on white matter in infants/children unexposed and living without HIV using DTI data^11–15^, only one has used tractography^13^. Unlike other approaches, tractography estimates white matter properties within a set of pairs of regions which provides anatomical context. It also allows for further study of the properties of connectivity between regions, using approaches such as graph analysis. While previous DTI studies report subtle exposure related regional differences in infants^15^ and children^11–14^, it isn’t yet known if these disrupt structural organization. In addition, no DTI study has yet included the possible influence of timing of maternal ART treatment initiation on developing white matter.

The work presented identifies potential HIV and ART exposure differences in white matter connections and networks shortly after birth in a cohort of HEU and HUU newborns. We proposed that maternal ART protects developing white matter from the effects of maternal infection. We hypothesized infants born to mothers on treatment at conception would demonstrate white matter integrity similar to HUU infants, whereas HEU newborns exposed to ART post conception would have altered regional connectivity. Lastly, we posited regional connectivity changes would affect network properties.

## Methods

### Study Cohort

The Healthy Baby Study (HBS) conducted in Cape Town, South Africa, included pregnant women living with and without HIV. The study enrolled 226 women, 18 years or older, at < 30 weeks’ gestation. The group comprised 82 women living without HIV and 144 women living with HIV. Among pregnant women living with HIV who were recruited, approximately half initiated ART prior to conception (*n*=78) and half initiated treatment post conception (*n*=66). As part of standard care in South Africa, HIV status is confirmed at the antenatal clinic using an HIV Rapid test. If positive, pregnant women living with HIV start first line therapy Tenofovir (TDF), Emtricitabine (FTC) and Efavirenz (EFV) in a daily fixed dose combination. HIV viral load is assessed at a follow up visit.

Exclusion criteria were underlying chronic disorders (e.g., diabetes, epilepsy, tuberculosis, hypertension), a history of recurrent premature deliveries, tuberculosis contact, use of medication other than essential pregnancy supplements, Isoniazid preventative therapy, or ART (if living with HIV), and for those with HIV, poor adherence to ART, not being on fixed drug combination ART (TDF, FTC, EFV), or nondisclosure of HIV status to family members. Prior to enrolment, mothers were also interviewed regarding their alcohol and drug use habits using the timeline follow back questionnaire^16,17^. Mothers engaged in illicit drug use, binge drinking (4 or more drinks per occasion) or drinking more than minimally (>7 drinks per week), were also excluded.

Pregnant women provided written informed consent for themselves and their infants to participate in the study. The HBS study was conducted in accordance with protocols approved by the Health Sciences Human Research Ethics Committees of Stellenbosch (ref: M16/10/041 and S21/11/231) and the University of Cape Town (ref: 801/2016).

Gestational age was determined by the study clinicians using date of Last Menstrual Period (LMP) and Ultrasound as per American College of Obstetrician and Gynaecologists (ACOG) (2017) guidelines^18^. If no ultrasound or accurate LMP was available, physical examination was used according to local guidelines^19^.

For mothers living with HIV, HIV viral load (VL), CD4 count and treatment records were obtained throughout pregnancy at antenatal visits.

For prevention of vertical transmission, infants born to women living with HIV were given Nevirapine if considered low risk. Infants at high risk of vertical transmission, defined as a maternal VL >1000 copies/mL at 32 weeks GA, are also prescribed Zidovudine. Infants were excluded if they were born earlier than 36 weeks GA, weighed less than 2500 grams at birth, received a positive HIV-1 PCR test, or were diagnosed with a condition that could influence neurodevelopment. Further details about the maternal and infant inclusion and exclusion criteria can be found in Ibrahim et al^20^.

Infants born to mothers living with HIV who initiated ART before conception are referred to as HEU pre-conception, or HEU_pre_. Similarly, neonates born to mothers living with HIV who started ART during pregnancy, are denoted as HEU post-conception, or HEU_post_.

### MRI acquisition

The study performed brain imaging on neonates using a 3T Siemens Skyra MRI scanner (Siemens, Erlangen, Germany) at the Cape Universities Body Imaging Centre (CUBIC) located at Groote Schuur Hospital. Prior to scanning, newborns had their diapers changed and were tightly swaddled to reduce motion. Sponge ear plugs and specially designed foam ear pads were placed in and over their ears, held in place by a beanie cap. The infants were fed and put to sleep in supine position on a special styrofoam bead pillow in a 16-channel paediatric head coil (Siemens), and imaged without sedation.

The protocol included a high-resolution T1-weighted (T1w) 3D echo-planar imaging (EPI) navigated multi-echo magnetization-prepared rapid gradient-echo (MEMPRAGE) acquisition (FOV 192×192 mm^2^, TR 2540 ms, TI 1450 ms, TEs = 1.69/3.55/5.41/7.27 ms, BW 650 Hz/px, 144 sagittal slices, voxel size 1.0×1.0×1.0 mm^3^). Two diffusion-weighted imaging (DWI) sets with opposite (Anterior-Posterior, Posterior-Anterior; AP/PA) phase encoding directions were acquired with a multi-band^21^ twice refocused spin-echo EPI sequence: TR 4800 ms, TE 84 ms, matrix 62 axial slices of 96×96 voxels (each voxel 2×2×2 mm^3^), 6/8 partial Fourier encoding, BW 1628 Hz/px, with slice-acceleration factor 2 and GRAPPA factor 2. Each acquisition contained six b = 0 s mm^-2^ (b0) reference scans and 30 DW gradient directions with b = 1000 s mm^-2^.

### Image processing

All analyses were carried out using a combination of predeveloped in-house scripts and tools available in standard software packages such as the Analysis of Functional Neuroimages (AFNI) toolbox^22^, the Tolerably Obsessive Registration and Tensor Optimisation Indolent Software Ensemble (TORTOISE) version 3.1.0^23^, Infant Freesurfer ^24^, and the Functional And Tractographic Connectivity Analysis Toolbox (FATCAT)^25^ within AFNI.

(AFNI)’s *fat_proc_imit2w_from_t1w* function was used to create T2-weighted (T2w) anatomical image imitation of the T1w structural image. Next, each subject’s DW images were quality checked across both AP and PA directions. Subjects with less than 15 (50%) viable diffusion directions (out of the 30 present) and/or no viable b0 images (out of the 6) were excluded from further analyses. After exclusions, the DIFF_PREP function in TORTOISE was used to correct for DWI artefacts such as motion and eddy current distortions. The DRBUDDI function combined the AP and PA directions to perform EPI distortion correction and created a final set of DW images for each subject^26^. AFNI’s *fat_proc_dwi_to_dt* function was used to estimate diffusion tensors and DTI parameters.

Regions of interest (ROIs) were extracted from T1w images using Infant Freesurfer version 4a14499, which uses an automated algorithm to segment the infant brain^24,27^. Infant FreeSurfer segments each hemisphere into 8 subcortical ROIs, lateral-, 3^rd^- and 4^th^ ventricles, cerebral and cerebellar cortex, and cerebral white matter, as well as 4 medial structures, namely vermis, midbrain, pons and medulla. The left and right hemispheric cerebral white matter were not included as seeds in our analyses. We included the ventricles because of their location relative to subcortical structures. The third ventricle is between the right and the left thalamus, while the fourth ventricle sits within the brainstem, at the intersection between the pons and medulla oblongata. The final set of DW images were co-registered to Tw1 images to map the seeds to the DWI space. Figure 1 shows the 28 ROIs that were used as seeds in our analyses.

**Figure 1:**
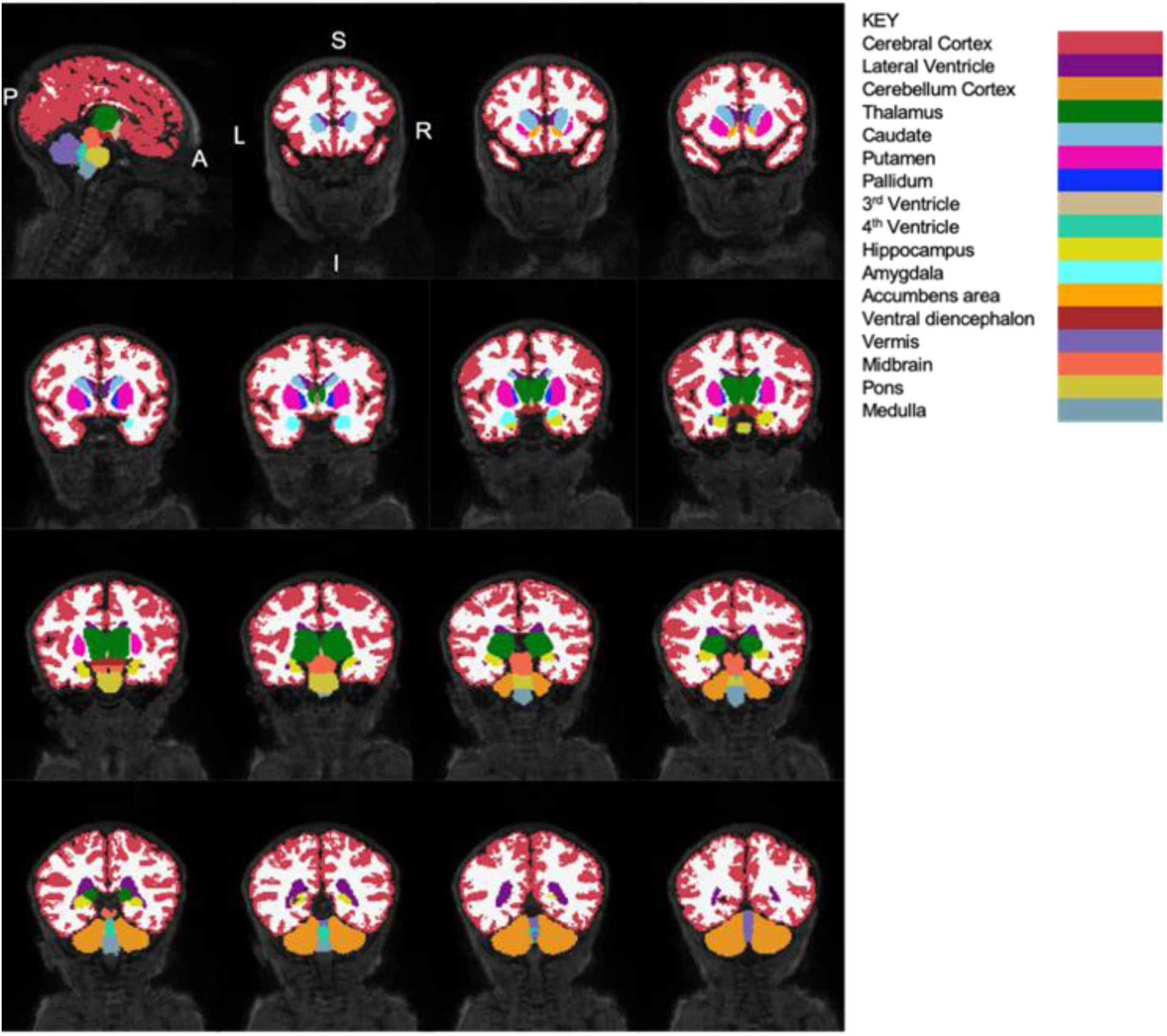
Sagittal and coronal views of neonatal brain (from a randomly selected HUU infant) showing the segmented structures and their corresponding labels (segmented in Infant Freesurfer). To visualise different structures, coronal slices are presented from the anterior to posterior direction. A/P – Anterior/Posterior; L/R – Left/Right; S/I – Superior/Inferior.

### DTI and Tractography

The diffusion metrics commonly extracted in DTI analysis include mean diffusivity (MD), fractional anisotropy (FA), axial diffusivity (AD) and radial diffusivity (RD). MD reflects the average amount of water diffusion within a voxel^28^. It is lower in regions of high tissue complexity as this creates diffusion obstacles^28^, and is typically low within white matter due to the large number of axonal connections present^28^. However, in infant brains, MD is often significantly higher than in mature adult brains because brain water content decreases during maturation^28^. In infant brains, the majority of axons are unmyelinated and structures such as cell and axonal membranes are less densely packed, so water is more readily able to diffuse perpendicularly. FA is used as an index of the amount of anisotropy within a specified region^29^. In tissue, FA describes the directional coherence of diffusivity and for this reason is used as a quantitative marker of white matter integrity^28^. The degree of anisotropy is associated with axon density and axon count, and while the degree of myelination relates to FA as well, it does not define tissue anisotropy.

AD, also known as longitudinal or parallel diffusion, refers to diffusivity on the principal axis of the diffusion ellipsoid. RD, also known as transverse or perpendicular diffusion, refers to diffusivity perpendicular to the principal direction of the diffusion ellipsoid, or an average of diffusion along the ellipsoid’s two minor axes^28^. Low AD is typically associated with axonal damage and fragmentation, while RD has been related to fiber coherence, myelin integrity, axonal diameter, and axonal density^28^. Presenting AD and RD can be helpful for interpretation, as changes in AD and RD typically drive changes in FA^30,31^.

Tractography was performed between every pair of seeds using the Functional And Tractographic Connectivity Analysis Toolbox (FATCAT) (version 1.1)^25^ in AFNI. The *3dTrackID* function was used to perform fully probabilistic tractography^32^. The stopping criteria for this study included the following: angle threshold (turning angle > 60°), FA threshold for infants (FA < 0.1) and tract length (<20 mm)^33,34^.

The outputs of the 3dTrackID are a set of the following measures – FA, MD, AD, RD, bundle length (BL), number of voxels in a bundle (NV), fractional volume of the bundle (fNV) and fractional number of streamlines in a given tract (fNT) – for each tract connecting the ROIs and subject. Due to inter-subject variability of data, sets of tracts for each subject could in principle be different. To ensure a common subset of tracts and DTI measures among the entire population in the study, we used the intersection of all the individual sets of tracts.

### Graph theory

Tractography parameters were used as inputs for graph theory analysis. We defined a graph as set of nodes (ROIs) and weighted edges (white matter tracts). The weight of each edge was defined by fNT.

Each subject’s structural n x n connectivity matrix (n being number of nodes) weighted by connections’ fNT gave rise to a weighted adjacency matrix for each subject. Graph analysis was performed using R statistical software^35^ and the igraph package for brain network analysis. The graph measures considered include strength, transitivity, local and nodal efficiency. A mean graph was created for each group (HUU, HEU_pre_ and HEU_post_) using the weighted adjacency matrices.

### Statistical analysis

Statistical analyses were performed using R statistical software^35^. We used linear regression models to compare the differences arising from HIV exposure and different ART durations (HUU vs HEU_pre_ and HEU_post_) in both DTI and graph theory measures.

DTI measures of interest were extracted, and outliers were removed from every common pairwise connection between target ROIs. Outliers were identified as values less than 1.5 times the interquartile range (IQR) below the lower quartile (Q_L_ - 1.5(IQR)) or 1.5 times the IQR above the upper quartile (Q_U_ + 1.5(IQR)). The outliers in the analyses of graph theoretical measures were removed in an analogous fashion.

DTI measures were then used in linear regression models. To identify confounders, we summarised the likely associations between potential confounders (infant sex, gestational age equivalent at scan, weight at scan, head circumference at scan, maternal weight gain per week of pregnancy, maternal education, ounces absolute alcohol consumed per day across pregnancy (oz AA/day) and maternal age at delivery), exposure variables and imaging outcome variables.

For the linear regression models, we chose a statistical significance level of p=0.05. Subsequently, we corrected the calculated p-values for multiple comparisons using the false discovery rate (FDR) method. The corrected p-values, presented as q-values, were deemed statistically significant at a level of q<0.05.

To explore the possible influence of maternal immune health on affected white matter integrity and network measures, we performed correlation analysis between DTI outcome measures and maternal CD4 count in pregnancy within tracts and networks showing group differences.

## Results

### Sample demographics

After exclusions, 187 infants were enrolled in the study (67 HUU, 63 HEU_pre_, 55 HEU_post_), of whom 185 visited CUBIC for neonatal MRI (2 HEU_pre_ infants missed their scan). Of the 160 neonates who had T1 and DTI images, 108 had complete sets of DTI scans (satisfying the quality check conditions outlined in the Methods section). A further two infants were excluded due to poor quality T1 images, leaving 106 infants for the analysis. The infant data presented consists of 35 HUU and 71 HEU (36 HEU_pre_ / 35 HEU_post_) infants. While the pre-conception HEU infant group was exposed to ART throughout gestation, the infants in the HEU_post_ group were exposed to ART *in utero* between 14 to 36 weeks. The sample demographics are presented in Table 1.

**Table 1:** Demographic data of HUU controls and HEU groups at birth and at time of MRI scanning. Mean and standard deviations (SD) presented as Mean ± SD.

|  | HIV unexposed<br>uninfected (HUU) | HIV exposed uninfected (HEU) |  |
| --- | --- | --- | --- |
|  |  | HEU<sub>pre</sub> | HEU<sub>post</sub> |
| Number of subjects (n) | 35 | 36 | 35 |
| Sex (Male: Female) | 18:17 | 18:18 | 20:15 |
| **Birth measures** |  |  |  |
| Gestational age (weeks) | 40 ± 1 | 40 ± 2 | 40 ± 2 |
| Weight (g) | 3285 ± 407 | 3215 ± 333 | 3250 ± 403 |
| Length (cm) | 50 ± 4 | 50 ± 2 | 50 ± 3 |
| ARV exposure length (weeks) | N/A | 40 ± 1 | 25 ± 6 |
| **MRI scan measures** |  |  |  |
| Age at scan (weeks) | 1.9 ± 1 | 1.5 ± 1 | 1.8 ± 1 |
| Gestational age equivalent (weeks) | 42 ± 1 | 41 ± 1 | 42 ± 1 |
| Weight (g) | 3561 ± 439 | 3143 ± 411 | 3509 ± 379 |
| Length (cm) | 52 ± 2 | 51 ± 2 | 52 ± 2 |
| Head circumference (cm) | 35 ± 1 | 36 ± 1 | 35 ± 1 |
| **Maternal measures** |  |  |  |
| Birth age (years) | 28 ± 6 | 31 ± 6 | 28 ± 5 |
| CD4 count within 6 months of pregnancy (cells/mm<sup>3</sup>) | N/A | 571 ± 180 | 435 ± 194 |
| *Distribution of peak Viral Loads (VLs) during pregnancy (n, %)* |  |  |  |
| <20 copies/mL | N/A | 25 (69%) | 18 (51%) |
| ≥ 20 – 400 copies/mL | N/A | 7 (19%) | 11 (31%) |
| ≥ 400 – 1000 copies/mL | N/A | 2 (6%) | 2 (6%) |
| ≥ 1000 copies/mL | N/A | 2 (6%) | 4 (11%) |
| Detectable VL throughout pregnancy (n, %) | N/A | 1 (3%) | 10 (29%) |
| Weight gain per week (g) | 0.41 ± 0.16 | 0.35 ± 0.28 | 0.27 ± 0.24 |
| Ounces absolute alcohol per day (oz AA/day)* | 0.02 ± 0.02 | 0.01 ± 0.02 | 0.02 ± 0.03 |

\*1 ounce AA ~ 2 standard drinks.

### DTI tractography

A total of 173 tracts were found through full-probabilistic DTI tractography to be common to all the subjects. We identified maternal weight gain per week of pregnancy and maternal education as confounding variables. Based on these tracts and confounders, we report the tracts showing HIV and ART exposure-related differences in FA and MD in Tables 2 – 4. Figures 2 and 3 represent the group differences across DTI measures using chord diagrams.

**Table 2:** Tracts showing lower fractional anisotropy (FA) in HEU_pre_ infants than HUU after FDR correction (q<0.05). Axial diffusivity (AD) and radial diffusivity (RD) in affected tracts are also shown. SD – standard deviation. L/R – Left/Right hemisphere.

| connections | FA |  |  |  |  |  | AD |  |  |  |  |  | RD |  |  |  |  |  |
| --- | --- | --- | --- | --- | --- | --- | --- | --- | --- | --- | --- | --- | --- | --- | --- | --- | --- | --- |
|  | mean HUU (SD) | mean HEU<sub>pre</sub> (SD) | p | q | std beta | std error | mean HUU (SD) | mean HEU<sub>pre</sub> (SD) | p | q | std beta | std error | mean HUU (SD) | mean HEU<sub>pre</sub> (SD) | p | q | std beta | std error |
| L CC - L Caud | 0.221 (0.007) | 0.214 (0.009) | 0.0006 | 0.036 | -0.479 | 0.137 | 1.633 (0.069) | 1.630 (0.044) | 0.990 | 0.990 | -0.002 | 0.016 | 1.172 (0.057) | 1.181 (0.031) | 0.379 | 0.456 | 0.126 | 0.132 |
| L CC - L Put | 0.208 (0.008) | 0.200 (0.008) | 0.0012 | 0.046 | -0.451 | 0.130 | 1.605 (0.063) | 1.618 (0.051) | 0.372 | 0.473 | 0.125 | 0.016 | 1.172 (0.060) | 1.194 (0.047) | 0.141 | 0.231 | 0.203 | 0.156 |
| L CC - L VDC | 0.224 (0.009) | 0.216 (0.008) | 0.0002 | 0.028 | -0.499 | 0.140 | 1.648 (0.061) | 1.653 (0.061) | 0.611 | 0.674 | 0.071 | 0.018 | 1.173 (0.058) | 1.191 (0.051) | 0.197 | 0.292 | 0.178 | 0.151 |
| L CC - R Thal | 0.235 (0.009) | 0.228 (0.011) | 0.0013 | 0.046 | -0.402 | 0.129 | 1.676 (0.078) | 1.705 (0.058) | 0.052 | 0.138 | 0.270 | 0.019 | 1.169 (0.054) | 1.197 (0.046) | 0.043 | 0.113 | 0.277 | 0.140 |
| L CC - MB | 0.225 (0.007) | 0.219 (0.007) | 0.0005 | 0.036 | -0.499 | 0.136 | 1.632 (0.062) | 1.647 (0.061) | 0.367 | 0.473 | 0.127 | 0.017 | 1.156 (0.050) | 1.182 (0.046) | 0.049 | 0.117 | 0.274 | 0.139 |

\*CC – Cerebral Cortex, Caud – Caudate, Put – Putamen, VDC – Ventral Diencephalon, Thal – Thalamus, MB - Midbrain

**Table 3:** Tracts showing higher mean diffusivity (MD) in infants in the HEU_pre_ group compared to those who are HUU after FDR correction (q<0.05). Axial diffusivity (AD) and radial diffusivity (RD) in affected tracts are also shown. SD – standard deviation. L/R – Left/Right hemisphere.

| connections | MD |  |  |  |  |  | AD |  |  |  |  |  | RD |  |  |  |  |  |
| --- | --- | --- | --- | --- | --- | --- | --- | --- | --- | --- | --- | --- | --- | --- | --- | --- | --- | --- |
|  | mean HUU (SD) | mean HEU<sub>pre</sub> (SD) | p | q | std beta | std error | mean HUU (SD) | mean HEU<sub>pre</sub> (SD) | p | q | std beta | std error | mean HUU (SD) | mean HEU<sub>pre</sub> (SD) | p | q | std beta | std error |
| L CC - R Caud | 1.306 (0.048) | 1.345 (0.057) | 0.0011 | 0.026 | 0.432 | 0.141 | 1.660 (0.041) | 1.703 (0.057) | 0.0004 | 0.017 | 0.475 | 0.014 | 1.134 (0.055) | 1.166 (0.060) | 0.0095 | 0.057 | 0.344 | 0.134 |
| L Thal - L Caud | 1.219 (0.043) | 1.247 (0.048) | 0.0019 | 0.032 | 0.414 | 0.128 | 1.519 (0.063) | 1.537 (0.059) | 0.0526 | 0.138 | 0.260 | 0.016 | 1.076 (0.044) | 1.099 (0.043) | 0.0107 | 0.057 | 0.342 | 0.128 |
| L Thal - L Put | 1.211 (0.048) | 1.242 (0.043) | 0.0043 | 0.046 | 0.381 | 0.129 | 1.524 (0.062) | 1.557 (0.054) | 0.0103 | 0.057 | 0.345 | 0.016 | 1.055 (0.043) | 1.081 (0.035) | 0.0072 | 0.050 | 0.364 | 0.129 |
| L Thal - L Amyg | 1.293 (0.044) | 1.345 (0.043) | 0.0000 | 0.001 | 0.616 | 0.136 | 1.602 (0.053) | 1.658 (0.058) | 0.0001 | 0.012 | 0.549 | 0.016 | 1.145 (0.045) | 1.189 (0.053) | 0.0006 | 0.027 | 0.473 | 0.146 |
| L Thal - L VDC | 1.248 (0.046) | 1.283 (0.041) | 0.0004 | 0.016 | 0.487 | 0.123 | 1.572 (0.057) | 1.607 (0.055) | 0.0022 | 0.023 | 0.427 | 0.016 | 1.087 (0.035) | 1.121 (0.041) | 0.0002 | 0.012 | 0.496 | 0.135 |
| L Thal - R VDC | 1.277 (0.050) | 1.303 (0.053) | 0.0033 | 0.046 | 0.388 | 0.133 | 1.632 (0.086) | 1.663 (0.063) | 0.0041 | 0.032 | 0.383 | 0.019 | 1.100 (0.045) | 1.126 (0.052) | 0.0047 | 0.047 | 0.376 | 0.137 |
| L Put - L Hipp | 1.297 (0.078) | 1.344 (0.046) | 0.0042 | 0.046 | 0.390 | 0.189 | 1.650 (0.095) | 1.694 (0.058) | 0.0095 | 0.055 | 0.353 | 0.021 | 1.120 (0.072) | 1.163 (0.052) | 0.0182 | 0.070 | 0.318 | 0.148 |
| L Hipp -L Amyg | 1.319 (0.060) | 1.389 (0.052) | 0.0001 | 0.004 | 0.519 | 0.151 | 1.617 (0.093) | 1.705 (0.077) | 0.0005 | 0.017 | 0.456 | 0.024 | 1.175 (0.057) | 1.230 (0.055) | 0.0012 | 0.035 | 0.414 | 0.146 |
| L Hipp - L VDC | 1.377 (0.076) | 1.429 (0.071) | 0.0043 | 0.046 | 0.396 | 0.149 | 1.719 (0.102) | 1.785 (0.068) | 0.0016 | 0.020 | 0.443 | 0.024 | 1.195 (0.081) | 1.250 (0.072) | 0.0027 | 0.043 | 0.409 | 0.129 |
| L Hipp - MB | 1.363 (0.073) | 1.416 (0.076) | 0.0013 | 0.026 | 0.429 | 0.132 | 1.713 (0.101) | 1.788 (0.105) | 0.0020 | 0.023 | 0.413 | 0.028 | 1.187 (0.067) | 1.231 (0.070) | 0.0030 | 0.044 | 0.394 | 0.138 |
| L VDC - R Thal | 1.267 (0.043) | 1.304 (0.043) | 0.0007 | 0.025 | 0.444 | 0.128 | 1.629 (0.063) | 1.668 (0.045) | 0.0040 | 0.032 | 0.401 | 0.015 | 1.087 (0.034) | 1.127 (0.041) | 0.0000 | 0.003 | 0.524 | 0.132 |
| L VDC - R VDC | 1.278 (0.043) | 1.319 (0.072) | 0.0010 | 0.026 | 0.434 | 0.161 | 1.637 (0.068) | 1.682 (0.073) | 0.0014 | 0.020 | 0.427 | 0.019 | 1.105 (0.052) | 1.130 (0.065) | 0.0236 | 0.081 | 0.299 | 0.144 |
| R CC - MB | 1.306 (0.046) | 1.341 (0.054) | 0.0047 | 0.047 | 0.384 | 0.142 | 1.620 (0.062) | 1.645 (0.059) | 0.0681 | 0.169 | 0.250 | 0.016 | 1.153 (0.045) | 1.190 (0.053) | 0.0041 | 0.047 | 0.385 | 0.147 |
| R LVent - MB | 1.321 (0.057) | 1.364 (0.055) | 0.0040 | 0.046 | 0.385 | 0.150 | 1.673 (0.073) | 1.731 (0.064) | 0.0005 | 0.017 | 0.451 | 0.018 | 1.145 (0.056) | 1.184 (0.054) | 0.0070 | 0.050 | 0.361 | 0.150 |
| R Thal -R Caud | 1.216 (0.042) | 1.256 (0.046) | 0.0001 | 0.004 | 0.521 | 0.127 | 1.510 (0.055) | 1.549 (0.055) | 0.0006 | 0.017 | 0.451 | 0.014 | 1.070 (0.029) | 1.106 (0.040) | 0.0000 | 0.001 | 0.584 | 0.139 |
| R Thal - R Hipp | 1.372 (0.056) | 1.406 (0.042) | 0.0035 | 0.046 | 0.424 | 0.145 | 1.695 (0.063) | 1.737 (0.051) | 0.0006 | 0.017 | 0.490 | 0.016 | 1.211 (0.056) | 1.240 (0.043) | 0.0163 | 0.069 | 0.353 | 0.142 |
| R Amyg-R VDC | 1.321 (0.059) | 1.359 (0.042) | 0.0014 | 0.026 | 0.426 | 0.148 | 1.654 (0.067) | 1.698 (0.055) | 0.0030 | 0.028 | 0.398 | 0.017 | 1.155 (0.056) | 1.190 (0.044) | 0.0021 | 0.041 | 0.404 | 0.136 |

\*CC – Cerebral Cortex, Caud – Caudate, Thal – Thalamus, Put – Putamen, Amyg – Amygdala, VDC – Ventral Diencephalon, Hipp – Hippocampus, MB – Midbrain, LVent – Lateral Ventricle

**Table 4:** Tracts where infants in the HEU_post_ group have lower (FDR q<0.05) fractional anisotropy (FA) than those who are HUU. Axial diffusivity (AD) and radial diffusivity (RD) in affected tracts are also shown. SD – standard deviation. L/R – Left/Right hemisphere.

|  | FA |  |  |  |  |  | AD |  |  |  |  |  | RD |  |  |  |  |  |
| --- | --- | --- | --- | --- | --- | --- | --- | --- | --- | --- | --- | --- | --- | --- | --- | --- | --- | --- |
| connections | mean HUU (SD) | mean $HEU_{post}$ (SD) | p | q | std beta | std error | mean HUU (SD) | mean $HEU_{post}$ (SD) | p | q | std beta | std error | mean HUU (SD) | mean $HEU_{post}$ (SD) | p | q | std beta | std error |
| L CC - L Thal | 0.209 (0.011) | 0.200 (0.010) | 0.0015 | 0.024 | -0.413 | 0.003 | 1.635 (0.067) | 1.652 (0.071) | 0.530 | 0.993 | 0.083 | 0.018 | 1.190 (0.061) | 1.218 (0.048) | 0.143 | 0.348 | 0.192 | 0.014 |
| L CC - L Caud | 0.221 (0.007) | 0.213 (0.011) | 0.0013 | 0.024 | -0.436 | 0.003 | 1.633 (0.069) | 1.646 (0.067) | 0.583 | 0.993 | 0.074 | 0.018 | 1.172 (0.057) | 1.188 (0.050) | 0.261 | 0.439 | 0.150 | 0.014 |
| L CC - L Put | 0.208 (0.008) | 0.198 (0.008) | 0.0001 | 0.010 | -0.542 | 0.002 | 1.605 (0.063) | 1.625 (0.065) | 0.392 | 0.993 | 0.114 | 0.017 | 1.172 (0.060) | 1.194 (0.056) | 0.174 | 0.365 | 0.177 | 0.015 |
| L CC - L Pal | 0.224 (0.009) | 0.217 (0.010) | 0.0066 | 0.048 | -0.372 | 0.003 | 1.620 (0.067) | 1.630 (0.063) | 0.638 | 0.993 | 0.063 | 0.017 | 1.152 (0.056) | 1.176 (0.059) | 0.123 | 0.328 | 0.201 | 0.015 |
| L CC - L Amyg | 0.220 (0.013) | 0.210 (0.013) | 0.0018 | 0.026 | -0.432 | 0.004 | 1.693 (0.058) | 1.709 (0.093) | 0.824 | 0.993 | 0.030 | 0.020 | 1.205 (0.054) | 1.236 (0.056) | 0.087 | 0.287 | 0.216 | 0.014 |
| L CC - L VDC | 0.224 (0.009) | 0.215 (0.006) | 0.0000 | 0.000 | -0.642 | 0.002 | 1.648 (0.069) | 1.658 (0.075) | 0.778 | 0.993 | 0.038 | 0.019 | 1.173 (0.058) | 1.192 (0.056) | 0.295 | 0.469 | 0.139 | 0.015 |
| L CC - R CC | 0.195 (0.008) | 0.189 (0.005) | 0.0075 | 0.048 | -0.372 | 0.002 | 1.620 (0.072) | 1.641 (0.054) | 0.286 | 0.993 | 0.140 | 0.017 | 1.208 (0.063) | 1.235 (0.047) | 0.038 | 0.265 | 0.272 | 0.014 |
| LCC - R LVent | 0.227 (0.009) | 0.221 (0.010) | 0.0057 | 0.048 | -0.376 | 0.002 | 1.742 (0.097) | 1.756 (0.090) | 0.809 | 0.993 | 0.032 | 0.025 | 1.227 (0.074) | 1.251 (0.064) | 0.306 | 0.477 | 0.139 | 0.019 |
| L CC - MB | 0.225 (0.007) | 0.215 (0.010) | 0.0007 | 0.023 | -0.474 | 0.003 | 1.632 (0.062) | 1.637 (0.070) | 0.918 | 0.993 | -0.014 | 0.018 | 1.156 (0.050) | 1.184 (0.045) | 0.061 | 0.274 | 0.246 | 0.013 |
| L CC - Pons | 0.224 (0.009) | 0.216 (0.010) | 0.0063 | 0.048 | -0.387 | 0.003 | 1.641 (0.067) | 1.652 (0.058) | 0.769 | 0.993 | 0.040 | 0.017 | 1.170 (0.055) | 1.199 (0.042) | 0.052 | 0.274 | 0.262 | 0.013 |
| L Cb - R CC | 0.179 (0.015) | 0.168 (0.010) | 0.0015 | 0.024 | -0.409 | 0.003 | 1.474 (0.079) | 1.476 (0.082) | 0.818 | 0.993 | 0.031 | 0.021 | 1.119 (0.066) | 1.152 (0.086) | 0.092 | 0.287 | 0.226 | 0.020 |
| L Cb - 4th Vent | 0.234 (0.011) | 0.223 (0.011) | 0.0011 | 0.023 | -0.457 | 0.003 | 1.596 (0.052) | 1.611 (0.065) | 0.662 | 0.993 | 0.058 | 0.015 | 1.106 (0.049) | 1.144 (0.052) | 0.008 | 0.250 | 0.339 | 0.013 |
| L Cb - R Cb | 0.174 (0.016) | 0.162 (0.009) | 0.0004 | 0.023 | -0.471 | 0.004 | 1.427 (0.050) | 1.424 (0.043) | 0.826 | 0.993 | -0.032 | 0.013 | 1.085 (0.058) | 1.111 (0.035) | 0.040 | 0.265 | 0.285 | 0.013 |
| L Cb - Pons | 0.185 (0.017) | 0.172 (0.015) | 0.0008 | 0.023 | -0.437 | 0.004 | 1.497 (0.073) | 1.522 (0.089) | 0.277 | 0.993 | 0.146 | 0.022 | 1.130 (0.068) | 1.174 (0.084) | 0.022 | 0.250 | 0.300 | 0.020 |
| L Cb - Medulla | 0.189 (0.019) | 0.176 (0.021) | 0.0021 | 0.028 | -0.401 | 0.005 | 1.501 (0.095) | 1.534 (0.105) | 0.201 | 0.979 | 0.170 | 0.026 | 1.125 (0.092) | 1.169 (0.092) | 0.058 | 0.274 | 0.251 | 0.024 |
| 4th Vent - R CC | 0.243 (0.020) | 0.231 (0.012) | 0.0078 | 0.048 | -0.355 | 0.005 | 1.595 (0.069) | 1.604 (0.047) | 0.542 | 0.993 | 0.081 | 0.015 | 1.100 (0.062) | 1.126 (0.040) | 0.062 | 0.274 | 0.244 | 0.014 |
| 4th Vent - Pons | 0.210 (0.024) | 0.198 (0.017) | 0.0078 | 0.048 | -0.357 | 0.006 | 1.570 (0.094) | 1.571 (0.097) | 0.973 | 0.993 | -0.004 | 0.025 | 1.139 (0.073) | 1.175 (0.084) | 0.080 | 0.287 | 0.228 | 0.020 |
| 4th Vent-Medulla | 0.193 (0.026) | 0.173 (0.019) | 0.0031 | 0.034 | -0.382 | 0.006 | 1.55 (0.110) | 1.583 (0.112) | 0.449 | 0.993 | 0.100 | 0.029 | 1.157 (0.094) | 1.220 (0.102) | 0.025 | 0.250 | 0.290 | 0.026 |
| R CC - R Cb | 0.201 (0.010) | 0.193 (0.011) | 0.0038 | 0.038 | -0.401 | 0.003 | 1.577 (0.053) | 1.577 (0.051) | 0.789 | 0.993 | -0.037 | 0.014 | 1.150 (0.049) | 1.176 (0.043) | 0.055 | 0.274 | 0.241 | 0.012 |
| R CC - R Thal | 0.207 (0.009) | 0.200 (0.009) | 0.0056 | 0.048 | -0.372 | 0.002 | 1.624 (0.063) | 1.634 (0.065) | 0.930 | 0.993 | 0.011 | 0.016 | 1.187 (0.058) | 1.203 (0.057) | 0.447 | 0.581 | 0.096 | 0.014 |
| R CC - R Put | 0.205 (0.011) | 0.198 (0.007) | 0.0067 | 0.048 | -0.355 | 0.002 | 1.589 (0.057) | 1.599 (0.056) | 0.723 | 0.993 | 0.047 | 0.015 | 1.161 (0.058) | 1.174 (0.055) | 0.433 | 0.571 | 0.101 | 0.014 |
| R CC - R VDC | 0.218 (0.013) | 0.210 (0.007) | 0.0074 | 0.048 | -0.356 | 0.003 | 1.640 (0.062) | 1.652 (0.055) | 0.615 | 0.993 | 0.069 | 0.016 | 1.169 (0.055) | 1.196 (0.058) | 0.153 | 0.361 | 0.189 | 0.015 |
| R Cb - Vermis | 0.168 (0.015) | 0.160 (0.008) | 0.0067 | 0.048 | -0.372 | 0.004 | 1.403 (0.067) | 1.398 (0.067) | 0.535 | 0.993 | -0.084 | 0.018 | 1.083 (0.052) | 1.090 (0.043) | 0.670 | 0.776 | 0.057 | 0.013 |
| R Cb - MB | 0.193 (0.017) | 0.185 (0.015) | 0.0073 | 0.048 | -0.357 | 0.004 | 1.475 (0.047) | 1.472 (0.071) | 0.408 | 0.993 | -0.111 | 0.016 | 1.097 (0.053) | 1.106 (0.044) | 0.669 | 0.776 | 0.057 | 0.013 |
| R Cb - Medulla | 0.188 (0.021) | 0.173 (0.013) | 0.0010 | 0.023 | -0.435 | 0.005 | 1.478 (0.077) | 1.501 (0.086) | 0.350 | 0.993 | 0.124 | 0.021 | 1.120 (0.084) | 1.149 (0.060) | 0.080 | 0.287 | 0.236 | 0.020 |
| R Put - Pons | 0.251 (0.012) | 0.240 (0.015) | 0.0026 | 0.032 | -0.422 | 0.004 | 1.574 (0.055) | 1.559 (0.058) | 0.251 | 0.993 | -0.162 | 0.016 | 1.076 (0.046) | 1.077 (0.040) | 0.937 | 0.959 | -0.011 | 0.012 |
| Vermis - Pons | 0.187 (0.016) | 0.177 (0.013) | 0.0029 | 0.033 | -0.392 | 0.004 | 1.484 (0.056) | 1.496 (0.090) | 0.952 | 0.993 | -0.008 | 0.020 | 1.108 (0.050) | 1.139 (0.066) | 0.070 | 0.274 | 0.232 | 0.015 |
| Vermis- Medulla | 0.186 (0.025) | 0.172 (0.013) | 0.0010 | 0.023 | -0.434 | 0.005 | 1.472 (0.103) | 1.483 (0.099) | 0.985 | 0.997 | -0.003 | 0.027 | 1.102 (0.086) | 1.134 (0.080) | 0.122 | 0.328 | 0.212 | 0.022 |

\*CC – Cerebral Cortex, Thal – Thalamus, Caud – Caudate, Put – Putamen, Pal – Pallidum, Amyg – Amygdala, VDC – Ventral Diencephalon, LVent – Lateral Ventricle, MB – Midbrain, Cb – Cerebellum, 4<sup>th</sup> Vent – 4<sup>th</sup> Ventricle

**Figure 2:**
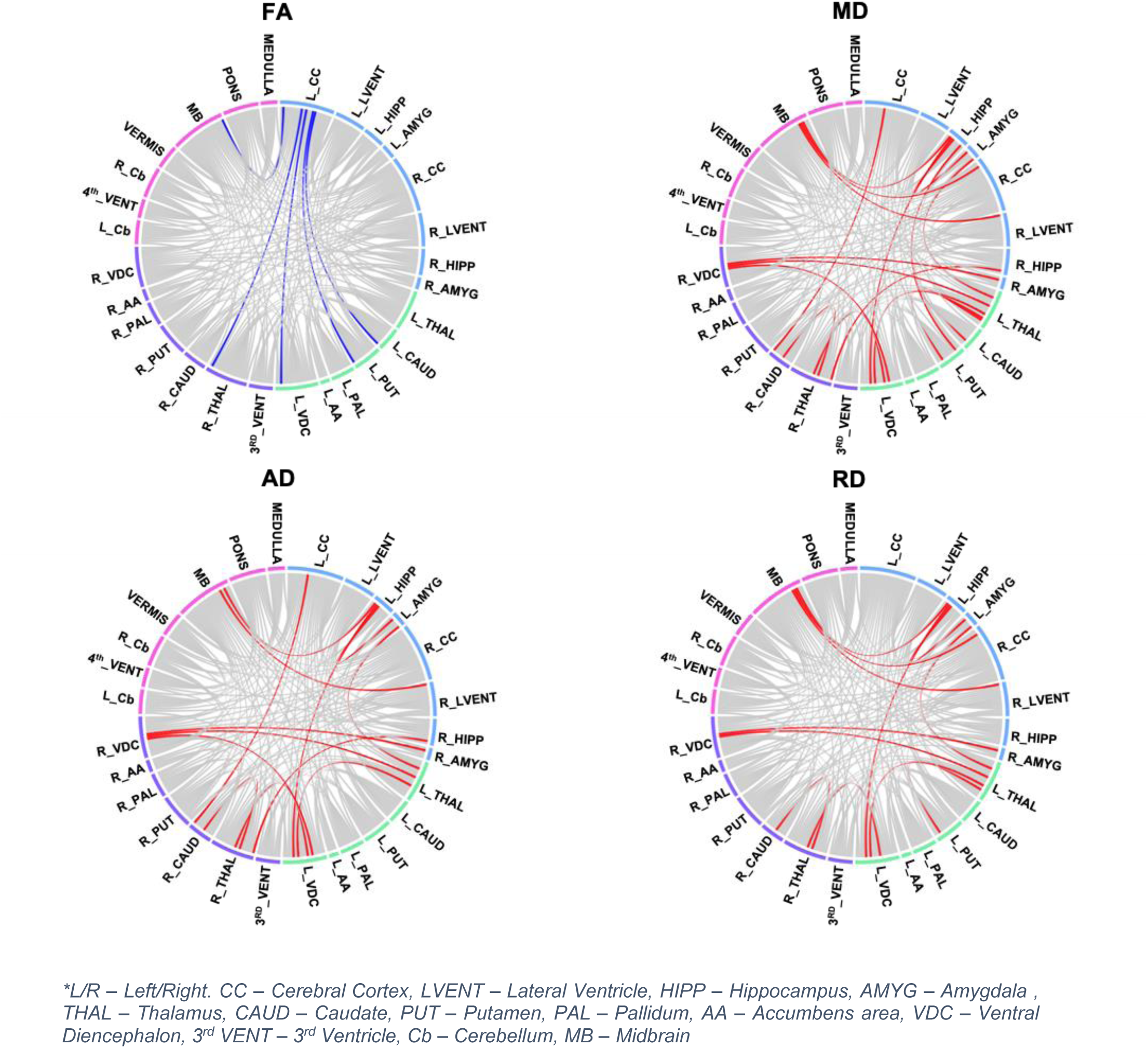
Chord diagram showing white matter connections with significant group differences in fractional anisotropy (FA), mean diffusivity (MD), axial diffusivity (AD) and radial diffusivity (RD) after FDR correction (q<0.05) between HUU and HEU_pre_ infants. The colours correspond to the communities present in the modularity graph, which include the left and right basal ganglia (in green and purple respectively), the forebrain (in blue) and the hindbrain (in pink).

**Figure 3:**
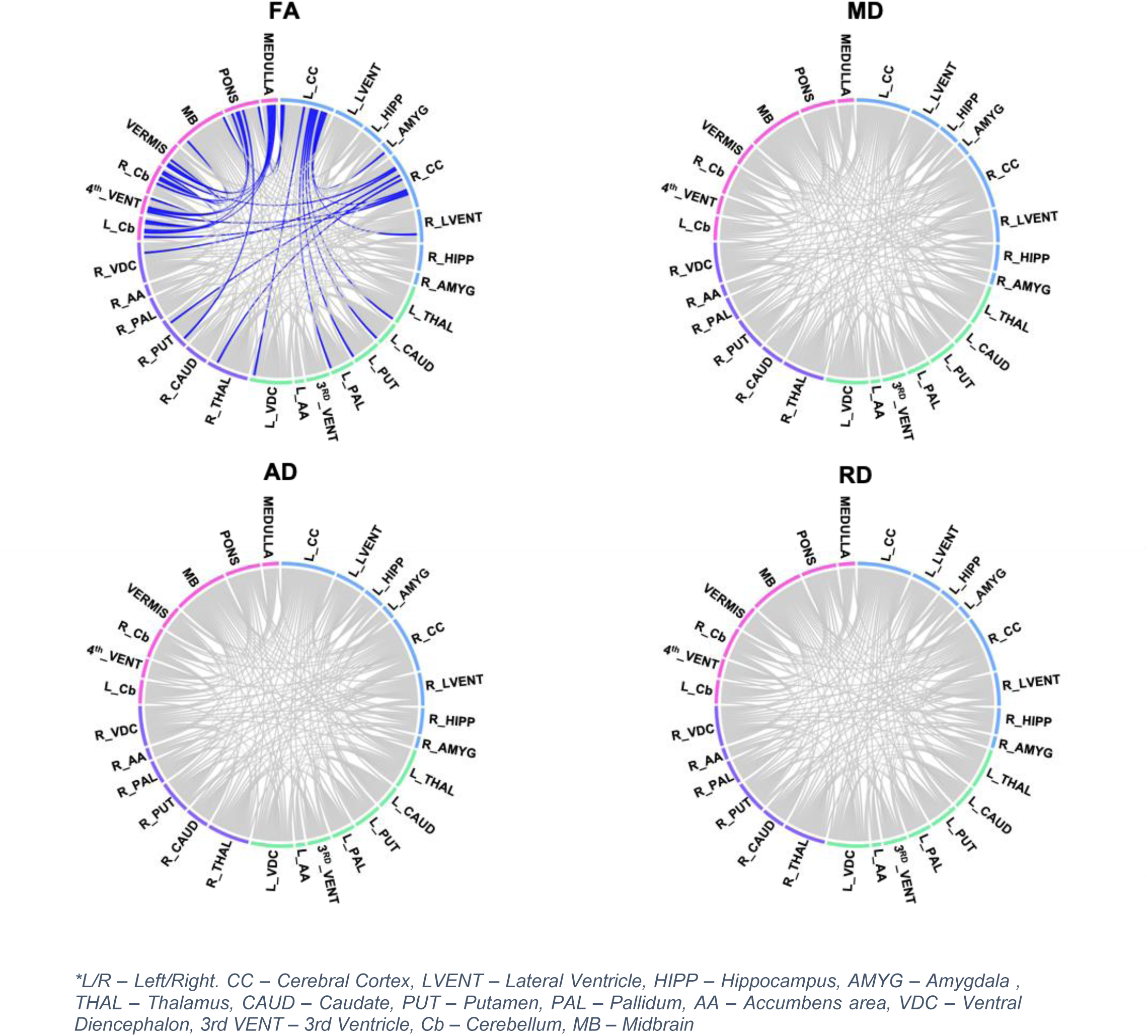
Chord diagram showing WM connections with significant group differences in fractional anisotropy (FA), mean diffusivity (MD), axial diffusivity (AD) and radial diffusivity (RD) after FDR correction (q<0.05) between HUU and HEU_post_ infants. The colours correspond to the communities present in the modularity graph, which include the left and right basal ganglia (in green and purple respectively), the forebrain (in blue) and the hindbrain (in pink).

Analysis shows reduced FA across five tracts between the left cerebral cortex and the midbrain, 1 right and 3 left subcortical structures in HEU infants exposed to ART since conception compared to HUU infants. We find lower mean FA in 28 white matter connections in HEU infants born to mothers who initiated ART post conception relative to HUU infants. All except 5 of these involve connections to either the cerebral cortex or the cerebellum. Notably, all 5 connections showing lower FA in HEU_pre_ infants, also demonstrate lower FA in the HEU_post_ group. While there are no significant group differences in AD or RD in the associated tracts after multiple comparisons, uncorrected p-values suggest FA reductions in at least a subset of these connections are likely due to increased RD.

In the HEU pre-conception group, we find seventeen tracts with increased MD as compared to HUU infants, 8 (47%) of which involve the thalamus. Higher MD in these tracts is accompanied by higher AD and RD.

Group analysis of bundle length, total number of tracts, number of voxels in a bundle and fractional volume of the bundle yielded no statistically significant group differences.

The HEU_pre_ group of neonates had significantly lower efficiency in the basal ganglia (left putamen and right accumbens area) compared to HUU infants after multiple comparison correction. In the analysis of HEU_post_ compared to HUU neonates, transitivity was significantly lower in the 4^th^ ventricle in the hindbrain. Results are summarized in Table 5.

**Table 5:** Graph theory measures with significant group differences after FDR correction (q<0.05).

| HUU vs HEU<sub>pre</sub> |  |  |  |  |  |  |
| --- | --- | --- | --- | --- | --- | --- |
| Nodal Efficiency |  |  |  |  |  |  |
| region name | mean HUU (SD) | mean HEU<sub>pre</sub> (SD) | p | q | std beta | std error |
| L Put | 2590 (2393) | 1186 (475) | 0.002 | 0.034 | -0.454 | 0.139 |
| R AA | 2697 (1903) | 2431 (2013) | 0.002 | 0.034 | -0.441 | 0.139 |
| HUU vs HEU<sub>post</sub> |  |  |  |  |  |  |
| Transitivity |  |  |  |  |  |  |
| region name | mean HUU (SD) | mean HEU<sub>post</sub> (SD) | p | q | std beta | std error |
| 4th Vent | 0.948 (0.012) | 0.939 (0.006) | 0.001 | 0.012 | -0.462 | 0.126 |

\*L/R – Left/Right, Put – Putamen, AA – Accumbens area, 4<sup>th</sup> Vent – 4<sup>th</sup> Ventricle

We found no significant associations between maternal CD4 and DTI based measures (FA, MD and graph outcomes) in any of the affected tracts.

## Discussion

Using DTI tractography, we identified neonatal white matter connections influenced by HIV and ART exposure duration in pregnancy in uninfected newborns. We found increased MD in white matter connections involving the thalamus and limbic system within HEU newborns who were exposed to ART since conception. Within the group of HEU neonates whose mothers initiated ART post conception, we observed reduced FA in cortical-subcortical as well as cerebellar connections. Overall, our analysis demonstrated distinct alterations in white matter integrity related to the timing of maternal ART initiation that influence local brain network properties.

### HIV exposure alterations

We found four tracts in the left hemisphere with lower mean FA values across HEU ART groups compared to HUU infants. These tracts connect the left caudate, putamen, ventral diencephalon and midbrain to the left cerebral cortex, reflecting projection fibers vulnerable to intrauterine HIV exposure independent of maternal ART. Projection fibers connecting cortical and subcortical regions begin development during the second trimester around 17 weeks gestational age, becoming more massive and voluminous between 18 and 30 post conception weeks^36^. Axonal fasciculation is the process in which previously formed axons provide guidance for growing axons, with axons adhering to one another. This occurs during the third trimester and leads to bundles of axons that follow similar growth patterns^37^. As this occurs, the developing white matter becomes more anisotropic, resulting in increased FA and AD, and decreased RD, while MD remains unchanged^10^. As such, these tracts with lower FA and higher RD (uncorrected p-value) may represent less organized localized projection fibers in HEU infants compared to their HUU peers.

Within the imaging literature of HEU pediatric populations, altered white matter properties in projection fibers have been observed previously. Using voxelwise methods, Jankiewicz et al. reports higher FA in 7-year-old HEU children in a cluster in the right corona radiata^11^, which connects the cerebral cortex to the thalamus and brainstem. While the direction of FA change and hemisphere are different than our findings, it is noteworthy that the same class of tracts is related to HIV-exposure in infancy and at 7-years.

Within this cohort, recently published work reported smaller left putamen volumes across all HEU infants, as well as smaller caudate volumes in infants whose mothers initiated ART post conception^20^. The reported growth disruption of these structures may contribute to the observed white matter alterations, however further work is needed to demonstrate statistical links between these findings.

### Pre-conception ART exposure alterations

We observed higher MD and RD/AD in HEU infants exposed to ART from conception compared to HUU infants, primarily in tracts that connect subcortical structures – only 2 of the 17 connections are to the cerebral cortex. Among the affected tracts, 8 involve the thalamus. The thalamus allows for communication between the body and brain^38^, relaying information and mediating the integration of information between specific cortical regions^39^. Once fiber bundles are organized pre-myelination occurs. During this period oligodendrocytes, glia cells that produce myelin, develop and mature. The process of oligodendroglial cell proliferation and development is isotropic, and is linked to decreased water and increased membrane density^40–42^. Taken together, this stage is characterized by constant FA and decreasing MD, RD and AD^10^. Our results in the pre-conception ART HEU group suggest pre-myelination in the affected tracts is hindered.

We find alterations in bilateral connections between the hippocampus, amygdala, thalamus and ventral diencephalon, which are part of the limbic system. These connections may belong to the fornix, a white matter component of the limbic system which connects the hippocampus to other subcortical structures in the limbic system. The fornix forms early, being visible in dissection studies by 13 weeks gestational age^43^.

In addition, we report higher MD in connections between the thalamus and striatum, specifically to the caudate and putamen. The altered connections may be thalamic subcortical projections in the thalamostriatal pathway, a part of the cortico-striato-thalamo-cortical circuit, which has been associated with regions in the frontal cortex involved in sensorimotor, limbic and cognitive information processing^39^.

In our cohort, the pre-conception HEU neonates were exposed to intrauterine ART during the first trimester as compared to the post-conception group. The most reported *in utero* ART effect on the brain is mitochondrial dysfunction^44^ and points to postponed pre-myelination in the affected tracts caused by decreased cellular metabolism^45^. Mitochondria are involved in creating cellular energy and mitochondrial disorder presents with a variety of neurological effects in infants and children. Our findings of higher MD in the pre-conception group suggest an ART effect on pre-myelinated white matter to/from the thalamus. However, further work is needed to link these alterations to mitochondrial dysfunction.

Among the pre-conception ART HEU group, graph analysis found lower nodal efficiency in the left putamen and right accumbens areas. Nodal efficiency is a measure of integration, estimating the ability to send information between a node and the network nodes^46,47^. The striatum plays an important role in facilitating information transfer between the cortex and thalamus. Our results suggest that altered subcortical white matter integrity, in the form of higher MD, AD, and RD, influences information processing in the basal ganglia community.

### Post-conception ART exposure alterations

In comparison to HUU newborns, connections with lower FA among newborns exposed to ART post-conception primarily involved cortical-subcortical as well as cerebellar connections. The cortical-subcortical tracts, like those also discussed in relation to the HIV-exposure, are projection fibers which begin development during the second trimester (13 to 28 weeks). This falls in the timeframe when mothers living with HIV in the post-conception group initiated treatment, 14 to 36 weeks post conception. The formation of axonal pathways takes place primarily after neuronal migration midway through pregnancy, and by birth at full term all major fibers are established^8^. As a result, growing axonal pathways are vulnerable during this period. The lower FA in these tracts may represent delayed axonal bundle organization due to maternal HIV disruptions to the fetal environment before ART was initiated.

Half of the connections with reduced FA in the post-conception ART HEU group involved the cerebellum and/or the brainstem. Coherent pathways of the cerebellar peduncles have been observed as early as 17 weeks gestational age^48^. Among voxelwise DTI studies in HEU infants and children, altered FA has been previously reported in the hindbrain. A study in newborns found increased FA in the cerebellar peduncles^15^, while a study in 10-year-old children reported decreased FA in the posterior cerebellum^12^. Taken together, our findings and the literature provide growing evidence that this region is vulnerable to maternal HIV exposure in infancy and early childhood.

Unlike the pre-conception ART exposure group, we observed reduced FA between the left and right cerebral cortex in infants exposed to ART *in utero* post conception. The pioneering axons of the corpus callosum, the largest set of commissural fibers, form around 13 gestational weeks^49,50^. From 13 to 23 weeks, fibers develop in an antero-posterior direction until the callosal fibers become uniform^43^. This window represents the period when mothers in this group were initiating ART. In the pre-conception group, ART may have protected the early development of the corpus callosum.

In terms of network measures, we observed lower transitivity around the 4^th^ ventricle of newborns whose mothers living with HIV initiated ART post conception as compared to the HUU group. The 4^th^ ventricle sits between the pons, medulla oblongata and cerebellum. The observed tracts involving the 4^th^ ventricle likely represent an adjacent structure(s) connected via the cerebellar peduncles. Transitivity is a measure of segregation related to specialized processing^47^. As compared to the HUU group, reduced transitivity may disrupt the development of specific domains within the interconnected hindbrain community structures. Considering the majority of tracts in the post-conception ART HEU group demonstrated lower FA in comparison to the HUU group in the hindbrain, these changes may contribute to the observed reduced transitivity.

### Data availability Statement

The datasets generated during and/or analysed during the current study are available from the corresponding author on reasonable request.

### Limitations

The white matter connections identified in this study are influenced by the size and number of seeds used for analysis. The Infant Freesurfer tool was used for segmentation, and the cortex is not subdivided into lobes or smaller divisions. As a result, the conclusions we can draw regarding connections to and within specific cortical regions are limited.

The DTI measures presented are derived from analysis based on several initial assumptions and are sensitive to methodological procedure^51^. Acquisition parameters, pre-processing steps, reconstruction/propagation models, and statistical analysis all affect the final sensitivity, specificity, and accuracy of a study^51,52^.

### Conclusions and Future Work

Using DTI tractography, we identified distinctive alterations in structural integrity and connectome measures in HEU infants related to maternal ART initiation. HEU newborns exposed to ART since conception demonstrated higher MD, RD and AD in connections between subcortical structures, many involving the thalamus. These changes suggest regional delayed pre-myelination. Further, reduced nodal efficiency in the striatum suggests altered subcortical white matter integrity influences the integration of information in the basal ganglia community. Within HEU newborns exposed to ART post-conception, we observed lower FA in cortical-subcortical and hindbrain connections, suggesting delays in axonal organization. In conjunction, we reported reduced transitivity in the hindbrain community which may influence regional functional processing. The initiation of ART before conception may have protected early white matter development in the hindbrain.

A recent meta-analysis of neurodevelopment outcomes in HEU infants found a risk of impairment in expressive language and gross motor development by age 2 years^5^. The predominant regions affected in both ART exposure groups, the thalamus, striatum, and cerebellum, are involved in aspects of language and motor development. The thalamus includes nuclei involved in motor activity, which relay and integrate information between the cortex and the basal ganglia or cerebellum^53^. Through the functional cortico-striato-thalamocortical neural pathways, the striatum is involved in the learning of motor skills^54^. The basal ganglia have been implicated in learning language, with basal ganglia abnormalities related to developmental disorders of language^55^. And, while it is well established that the cerebellum is involved in motor function, it also plays a role in various brain functions such as language, visuospatial ability and attention^56–59^. As a result, the reported abnormalities may contribute to later neurodevelopmental delays in language and motor domains in these infants. Further work is needed to explore whether our findings at the white matter connection or network level contribute to cognitive deficits in this cohort.

## Author Contributions Statement

NM: data analysis, interpretation of results, figure creation, drafting and review of manuscript

MH: oversight, project design, data analysis, interpretation of results, drafting and review of manuscript.

EM: conception and design of study, study oversight, data acquisition, review of manuscript.

FLW: data curation and review of manuscript

FL: review of manuscript

AvdK: conception and design of study, study oversight, data acquisition and review of manuscript.

BL: conception and design of study, study oversight, data acquisition and curation, and review of manuscript.

MJ: oversight, project design, data analysis, interpretation of results, and review of manuscript.

